# Hypomorphism of a novel long ERα isoform causes severe reproductive dysfunctions in female mice

**DOI:** 10.1101/2022.03.22.485385

**Authors:** Kenji Saito, Jacob E. Dickey, Samuel R. Rodeghiero, Brandon A. Toth, Matthew J. Kelly, Yue Deng, Uday Singh, Guorui Deng, Jingwei Jiang, Huxing Cui

## Abstract

Estrogen receptor alpha (ERα)-mediated estrogen signaling play a pivotal role in both reproductive and non-reproductive functions. Transcriptional regulation of ERα gene is highly complex, with multiple transcript variants being differentially produced across the tissues. However, tissue-specific variation and physiological specificity of the ERα variants are not yet fully understood. In an attempt to generate a Cre-dependently restorable ERα-null mice for functional investigation of the genetic sufficiency, we unexpectedly produced ERα hypomorphic mice with biased downregulation of a previously unappreciated long ERα isoform that is enriched in the female reproductive organs (uterus and ovaries) and the pituitary but minimally expressed in the brain. Female homozygous mutant mice were capable of pregnancy but displayed irregular estrus cycle and rarely maintained alive newborns without significant morphological and pathological changes in reproductive system and the disruption of body weight homeostasis, indicating the vital role of this long isoform in female reproductive function. Collectively, our results define a tissue-specifically enriched long ERα isoform and its preferential role in female reproductive function over body weight homeostasis.

## Introduction

Estrogens are steroid hormones known to regulate a wide range of physiological functions, including but not limited to reproduction, cardiovascular physiology, homeostatic regulation of energy balance, as well as a variety of social and learning behaviors(1). Estrogenic signaling is mediated through its binding to the cognate receptors, including estrogen receptor α, β (ERα/β), and a G protein-coupled estrogen receptor (GPR30)(2). Among them, emerging evidence indicates that ERα signaling has different impacts on different tissues or cells, leading to diverse physiological outcomes. Genetic studies in mice have revealed that ERα is a critical estrogen receptor for different aspects of reproductive function and energy homeostasis(3-7).

ERα belongs to a nuclear receptor superfamily of ligand-inducible transcription factors. The overall structure of the nuclear receptors is well conserved among the different subfamilies, which includes a N-terminal transactivation domain, a DNA-binding domain, a ligand-binding domain, and a C-terminal transactivation domain(8). The expression of ERα is highly regulated at transcriptional, post-transcriptional, and post-translational levels(9). Recent studies have revealed a highly complex nature of transcriptional regulation of ERα gene. Multiple ERα transcript variants are differentially produced among the tissues by transcription starts from different leader exons and alternative splicing(10-20). Alternative promoter usage and alternative exon splicing are known to occur in the mouse, rat, and human(10, 15, 16, 19, 21, 22), indicating that the complexity of ERα gene is conserved among species. However, the physiological significance of this complex transcriptional regulation is largely unexplored. Despite the presence of the different transcripts, so-called 66-kDa full-length ERα (ERα66) is synthesized from multiple transcript sequences(16). In addition to the leader exons, alternative splicing of internal exons generates numerous splice variants with either distinct 5’UTR or the proteins missing different functional domains(8). Some of these splice variants have either constitutive activity, no activity, or inhibitory effect on reference ERα(23). This complex ERα gene expression system is further complicated by post-translational modifications including phosphorylation, acetylation, sumoylation, ubiquitination, glycosylation, and palmitoylation(24, 25). However, it is worthy to note that most of functional studies have been performed without differentiating ERα splice variants and therefore, it remains largely unknown how these ERα variants differentially mediate the physiological and pathological roles of E2(8). A few exceptions include genetic mouse models of post-translation modifications in a phosphorylation site or palmitoylation site of ERα(26, 27). Since exons missing in a certain transcript variant are shared by other splice variants and genomic *cis* elements regulating splice variant expressions are largely unknown, *in vivo* genetic studies for specific physiological functions of splice variants have been extremely difficult.

Here we attempted to generate a Cre-dependently restorable ERα-null mouse model by inserting a transcription blocking (TB) cassette in ERα gene. Unexpectedly, our gene targeting strategy resulted in biased knockdown of specific long ERα isoform with molecular weight slightly higher than ERα66 (termed ERα66* herein), which enabled us to study the variant-specific physiological functions. Homozygous female mice carrying these hypomorphic alleles showed subfertility (instead of full infertility seen in previous ERα-null mouse models) without affecting body weight. We further show that this long ERα66* isoform is enriched in organs important for female reproductive function, including the ovary, testis, and pituitary, but minimally expression in the brain. Our findings shed new light on a previously unappreciated long ERα isoform and its vital role in female reproductive physiology.

## Results

### Generation and validation of mouse model

ERα, or Esr1, is expressed in both reproductive and non-reproductive organs where it meditates estrogenic signaling for a variety of different physiological functions. Germline deletion of ERα in mice causes infertility(4, 5), increased body weight(3), and various behavioral deficits(28, 29). To address the functional sufficiency of tissue-specific role of ERα, we attempted to generate a mouse model that enables conditional ERα restoration from the null background using the CRISPR-Cas9 system. Similar to previous studies (30-33), the deletion of ERα gene was attempted by the insertion of a loxP-flanked TB cassette into a putative intron located upstream of the first coding exon of ERα gene (ENSMUSE00001204758, Figure 1A). The TB cassette includes the following elements in order: a splice acceptor (SA) site from the mouse engrailed 2 gene (En-2) followed by green fluorescent protein with N-terminal fusion of the cytochrome c oxidase subunit VIII mitochondrial targeting signal (mitoGFP), an SV40 poly(A) signal, an SV40 enhancer, a synthetic poly(A) signal/transcriptional pause signal, and another synthetic poly(A) signal followed by a Myc-associated zinc finger protein-binding site. We took this transgenic strategy in order to enable Cre recombinase-mediated tissue-specific restoration of ERα expression for future experiments. We used mitoGFP instead of conventional GFP with the hope to study ERα bound to mitochondrial DNA in future experiment (34, 35). This feature enables immunocapture of mitochondria and *in vivo* or *ex vivo* imaging of mitochondrial dynamics within ERα-expressing cells. Genomic DNA was extracted from homozygous mutant mouse (hereafter referred as G/G), and PCR and sequencing of ERα genomic region confirmed the expected genotypes (Figure 1B) and correct CRISPR insertion (data not shown). Heterozygous (+/G) intercrossing produced wild-type (WT or +/+), +/G and G/G mice with expected Mendelian ratio (Table S1), indicating that gametes bearing only mutated G allele can proceed fertilization.

**Figure 1.**
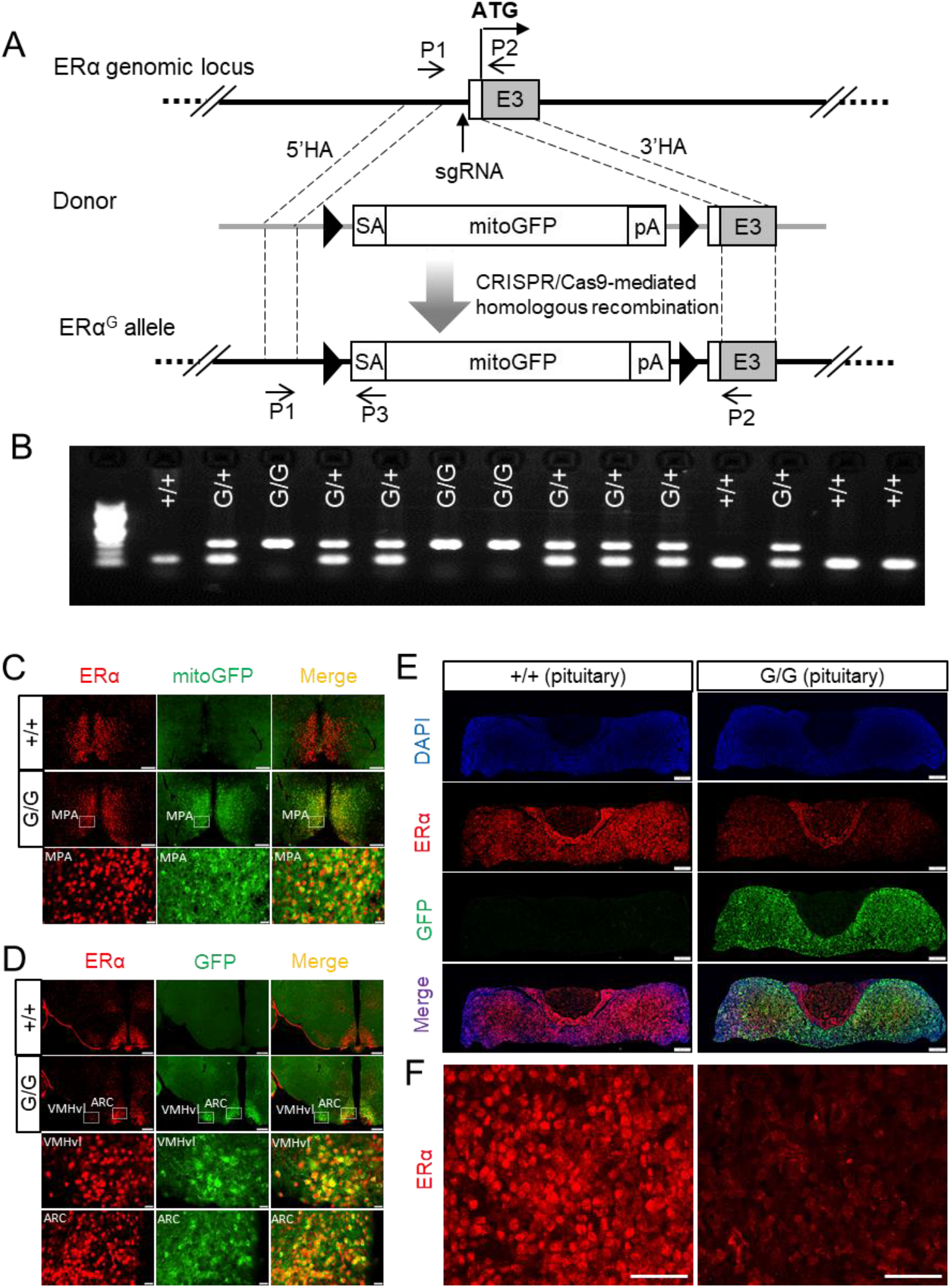
Generation and validation of ERα mutant (G/G) mouse. (A) Schematic showing the targeting strategy for the introduction of transcription blocking cassette consists of splicing acceptor (SA), mitochondrially targeted GFP (mtGFP), and polyadenylation signal (pA), into 5’ upstream of mouse ERα first coding exon 3 by CRISPR-Cas9-mediated homologous recombination. P1, P2, and P3 denote genotyping primers. HA, homology arm; pA, polyadenylation signal. (B) Representative gel image showing the genotyping results. (C-E) Representative images showing ERα and GFP immunoreactivity in the medial preoptic area (C), the ventrolateral division of VMH and the arcuate nucleus of hypothalamus (ARC) (D), and the pituitary (E) of +/+ and G/G female mice. (F) Zoom-in confocal image showing ERα immunoreactivity in the pituitary from E. Scale bar, 200 µm for C-E; 50 µm for F.

ERα is widely expressed throughout the brain with higher expression levels locally in some nuclei, such as medial preoptic area (MPA), medial amygdala, ventromedial (VMH) and arcuate (ARC) nuclei of hypothalamus (36, 37), and neuronal ERα plays critical roles in regulating energy and glucose homeostasis(6, 7). We therefore first examined GFP and ERα immunoreactivity in the brain sections of WT (+/+) and G/G mice by fluorescent immunohistochemistry (FIHC). We observed strong GFP expression in the MPA, ventrolateral division of VMH (VMHvl) and ARC of G/G, but not WT, mice (Figure 1C, D). GFP immunoreactivity was also detected in various other brain regions where endogenous ERα immunoreactivity has been reported, including the bed nucleus of stria terminalis, medial amygdala, periaqueductal gray, parabrachial nucleus, and spinal cord (data not shown). To our surprise, however, endogenous ERα immunoreactivity was also normally detected even in the brains of G/G mice, which nicely overlapped with GFP signals (Figure 1C, D), indicating the inability of inserted TB cassette to block endogenous ERα expression in the brain. ERα is also abundantly expressed in the pituitary and involves in the feedback regulation on the gonadotropin release (38-40). FIHC in the pituitary revealed a strong GFP immunoreactivity in G/G, but not WT, mice. Interestingly, however, in large contrast to normal ERα immunoreactivity in G/G brains, pituitary ERα was significantly reduced in G/G mice compared to WT littermates (Figure 1E, F). Taken together, these results indicate that our knockout strategy with inserted TB cassette works partially in the pituitary, but not the brain.

### Impaired reproductive function, but not energy balance, in G/G mice

Given the reported body weight gain in ERα KO mice(3, 41), we monitored body weight for both WT and G/G mutant mice. Consistent with normal expression of ERα in the brain, we did not observe significant body weight gain in G/G mutant mice compared to WT littermates at both young and older ages (Supplemental figure S1A-D). There was a trend toward increased body weight in 5-month-old G/G female mice compared to WT littermates but did not reach statistical significance (Supplemental figure S1C). ERα expression in the pituitary is a part of hypothalamus-pituitary-gonad (HPG) axis to affect female reproductive functions. We therefore further evaluated reproductive phenotypes of G/G female mice. Examination of the estrous cycle stage for consecutive eleven days (taken in the early afternoon) by vaginal cytology revealed that, in contrast to normal periodic transitioning of all estrus stages in WT female mice, G/G female mice rarely entered to estrus stage (Figure 2A). Irregular estrus cycle in G/G female mice prompted us to further evaluate their fertility. To evaluate female fertility, control (+/+ and +/G) and G/G female mice were mated with fertility-proven control male mice. Eight out of nine control females showed pregnancy and produced 50 alive newborns (6.25/liter on average) (Figure 2B). Surprisingly, seven out of eight G/G female also showed pregnancy (Figure 2B), indicating that G/G female mice likely can maintain normal sexual receptivity, ovulation, conception, and implantation. However, pregnant G/G female mice rarely delivered alive newborns (Figure 2B); all but one newborn was found dead in pregnant G/G female mouse cages. To evaluate whether neonatal death was preterm or postterm, we videotaped a few labors of pregnant G/G female mice. In a successful observation, we confirmed that a newborn was alive immediately after the delivery, but G/G mother did not take care of newborns after the labor (data not shown), suggesting that G/G female mice may have a defect in maternal behavior after the delivery. Additionally, we observed a significant reduction in uterine weight in G/G female mice compared to WT littermates upon tissue collection (Figure 2C). Given the reported abnormal histology of female organs in germline ERα knockout mice(5, 41, 42), we also performed standard hematoxylin and eosin (HE) staining on female reproductive tracts (vagina, uterus, and ovaries) in three +/+, two +/G, and three G/G female mice and found no obvious histological changes in G/G female mice compared to control littermates (Supplemental figure S2A-F). ERα Western blot (WB) analysis in the uterus revealed that, in contrast to clear multiple bands in a reference WT brain sample (denoted as #), the strong smear bands around 50-70 kDa were observed in WT uteri (Figure 2D) and the quantification of these signals confirmed more than 50% reduction of ERα protein level in G/G females compared to WT littermates (Figure 2E).

**Figure 2.**
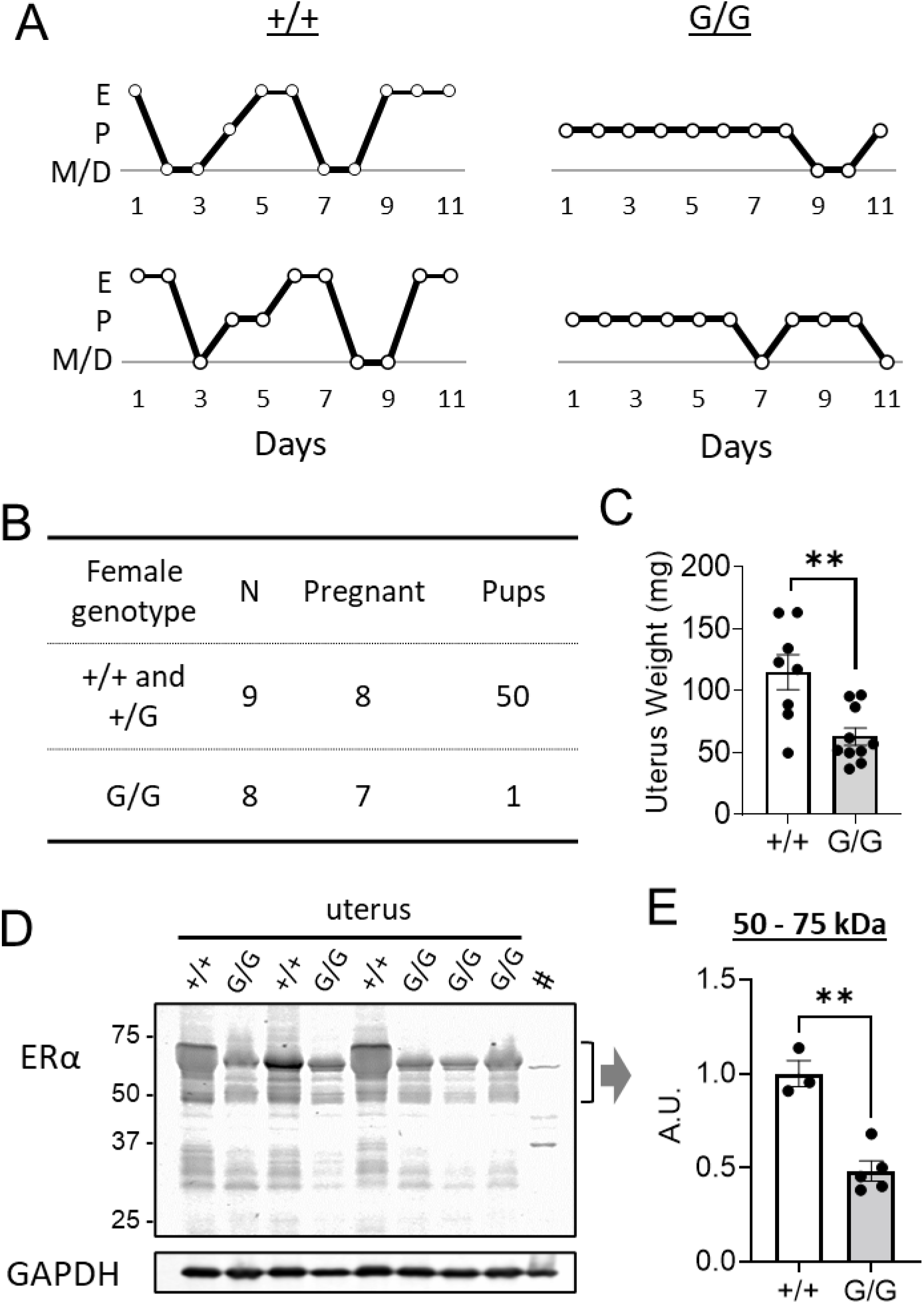
Reproductive phenotype in female G/G mice. (A) Representative traces of estrous cycle in two pairs of +/+ and G/G female mice. (B) A table summarizing the results of fertility test in +/+, +/G, and G/G female mice. (C) Uterus weight upon tissue collection. (D) The image of ERα WB in the uterus. Note that # denotes a reference brain sample that we used for ERα WB in different tissues in this study. (E) Quantification of 50-70 kDa ERα band shown in D. The data are expressed as means ± SEM. **p < 0.01 by unpaired student’s t-test.

### Biased knockdown of a novel long ERα66* isoform across the tissues

Although FIHC showed an unexpected normal retention of endogenous ERα while expressing GFP in the G/G brains, the TB cassette efficiently knocked down, but not knocked out, ERα in the pituitary and the uterus. Several ERα isoforms with different molecular weights have been reported(8, 43, 44). The smear bands around 50-70 kDa in WT uterus samples may reflect such isoforms and the TB cassette may differentially affect them. Therefore, we performed WB in other ERα-rich organs to evaluate whether the expression of ERα variants is differentially affected by the insertion of TB cassette in those tissues. As implied by FIHC, ERα protein level in the brain did not show remarkable genetic difference in ERα isoforms, including 66 kDa and other short bands we denoted as B-1, B-2, and B3 here (Figure 3A, C). However, we noticed an additional weak band on top of 66 kDa (denoted as 66* kDa) and the quantification revealed that only this 66* kDa band, but not 66 kDa and B-1-3, was specifically reduced in both male and female G/G mice (Figure 3B, D).

**Figure 3.**
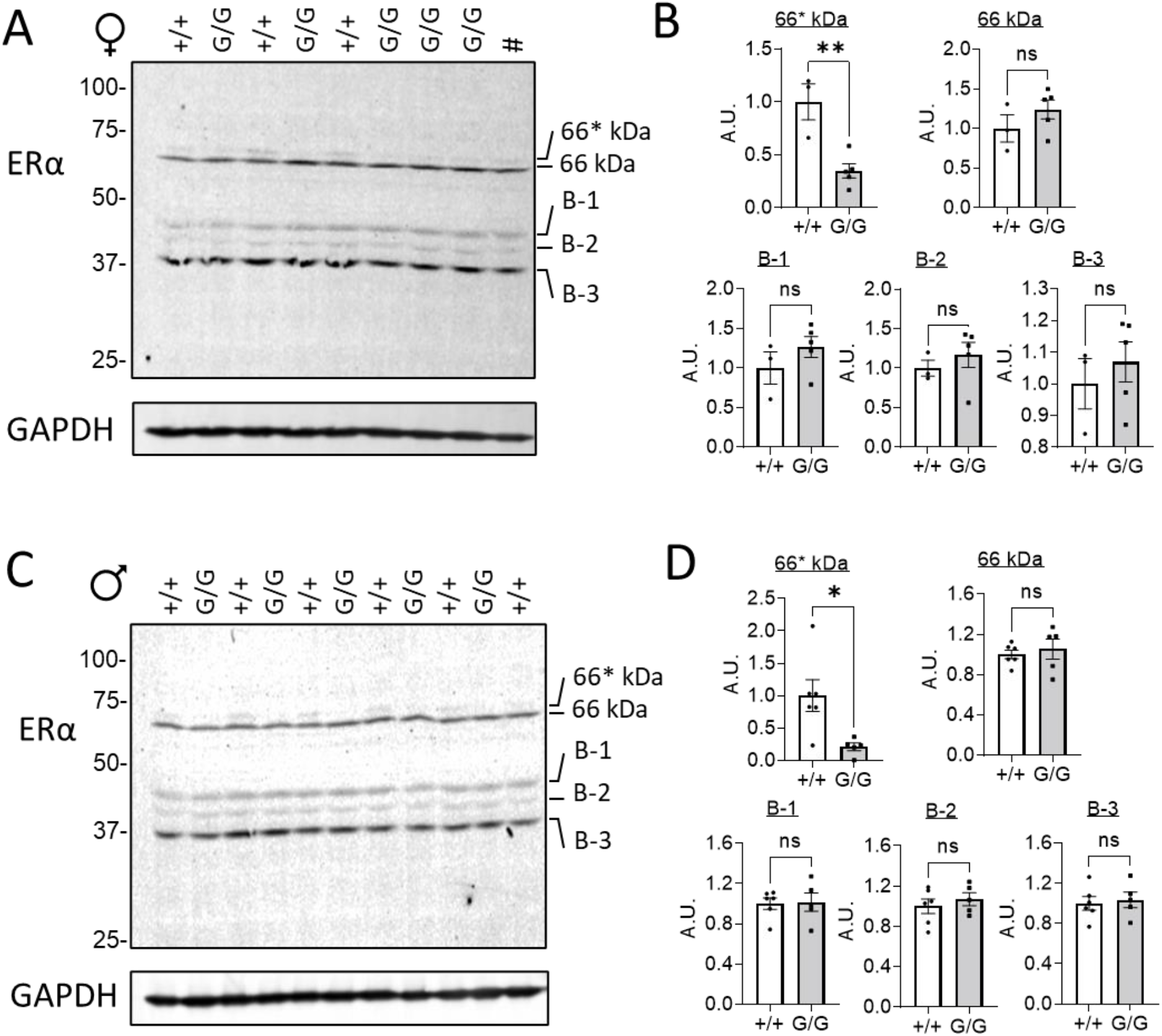
The ERα WB analysis in the brains of both male and female +/+ and G/G mice. (A) ERα and GAPDH (loading control) WB images in the brain of female mice (# denote a reference brain sample). (B) Quantification of different ERα bands shown in A. (C) ERα and GAPDH (loading control) WB images in the brain of male mice. (D) Quantification of different ERα bands shown in C. The data are expressed as means ± SEM. *p < 0.05, **p < 0.01, by unpaired student’s t-test.

ERα WB in the pituitary along with a reference brain sample (#) revealed that, in sharp contrast to dominant 66 kDa and other 3 B-1-3 bands in the brain, top 66* kDa band is much more abundant than 66 kDa in the pituitary without noticeable other short isoforms (Figure 4A, C). The quantification confirmed more than 50% reduction in 66* kDa, but not 66 kDa, in both female and male G/G mice compared to WT littermates (Figure 4B, D).

**Figure 4.**
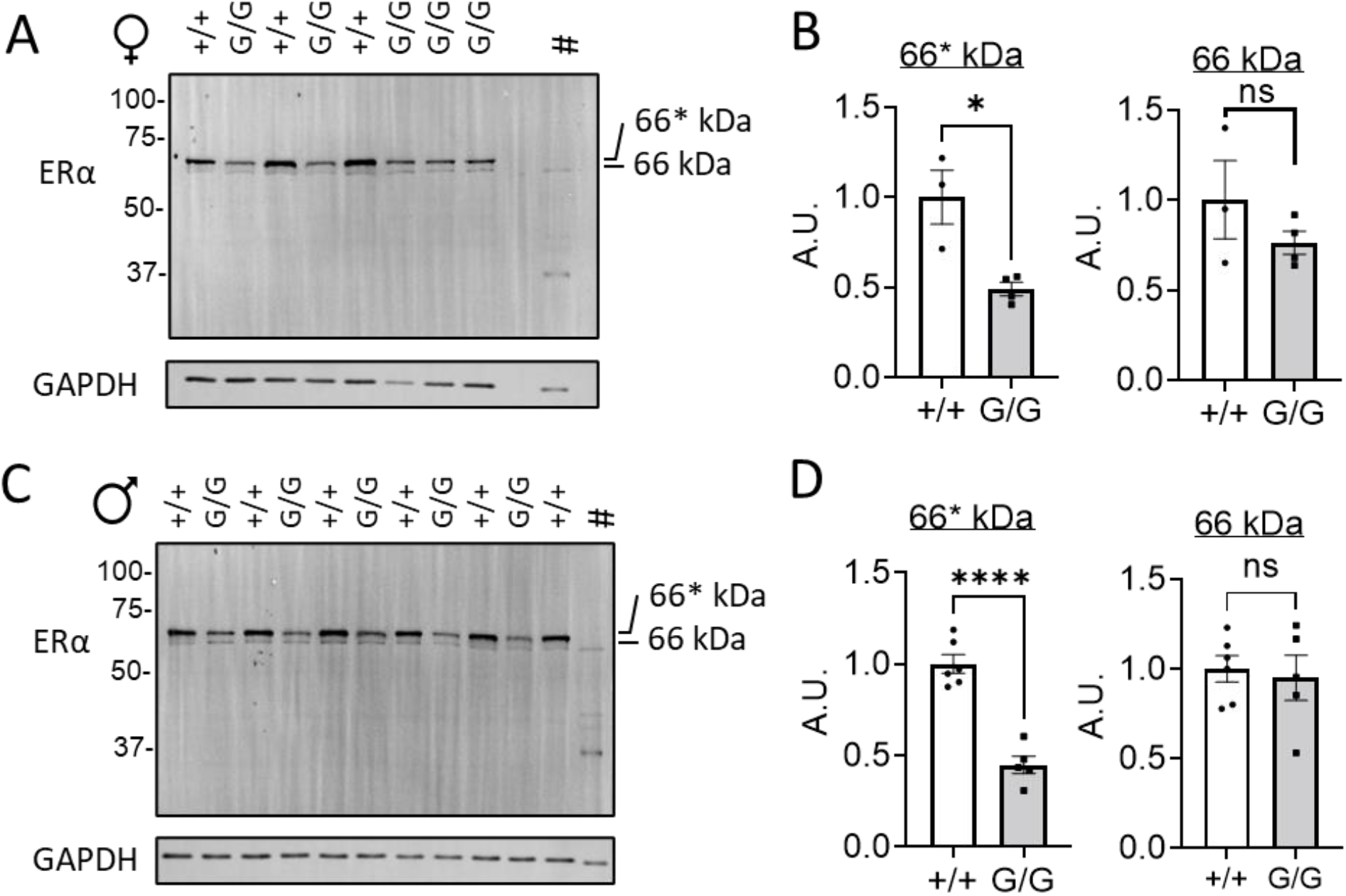
The ERα WB analysis in the pituitary of both male and female +/+ and G/G mice. (A) ERα and GAPDH (loading control) WB images in the pituitary of female mice (# denote a reference brain sample). (B) Quantification of 66* kDa and 66 kDa ERα bands shown in A. (C) ERα and GAPDH (loading control) WB images in the pituitary of male mice (# denote a reference brain sample). (D) Quantification of 66* kDa and 66 kDa ERα bands shown in C. The data are expressed as means ± SEM. *p < 0.05, ****p < 0.0001, by unpaired student’s t-test.

In ovaries, we found that both 66* kDa and 66 kDa ERα isoforms are highly expressed, although long 66* kDa appears to be the dominant (Figure 5A). We also observed a subdominant band around 50 kDa (denoted as OV-1; Figure 5A) that was not observed in the brain and the pituitary. The quantification revealed more than 60% reduction of 66* kDa and OV-1, but not 66 kDa, isoforms in G/G females compared to WT littermates (Figure 5B). In sharp contrast to the ovaries, 66 kDa appears to be the dominant over 66* kDa in the testes and there was another major band around 30 kDa (denoted as TE-1; Figure 5C). The quantification revealed a trend toward reduced 66* kDa, but not 66 kDa and TE-1, in G/G males compared to WT littermates but did not reach a statistical significance (Figure 5D). Interestingly, OV-1 (∼50 kDa) band observed in female ovaries was absent in male testes. These results indicate that ERα isoforms are differentially expressed across the tissues and mutant G/G mice have specific knockdown (>50%) of 66* kDa isoform across different endocrine and reproductive organs.

**Figure 5.**
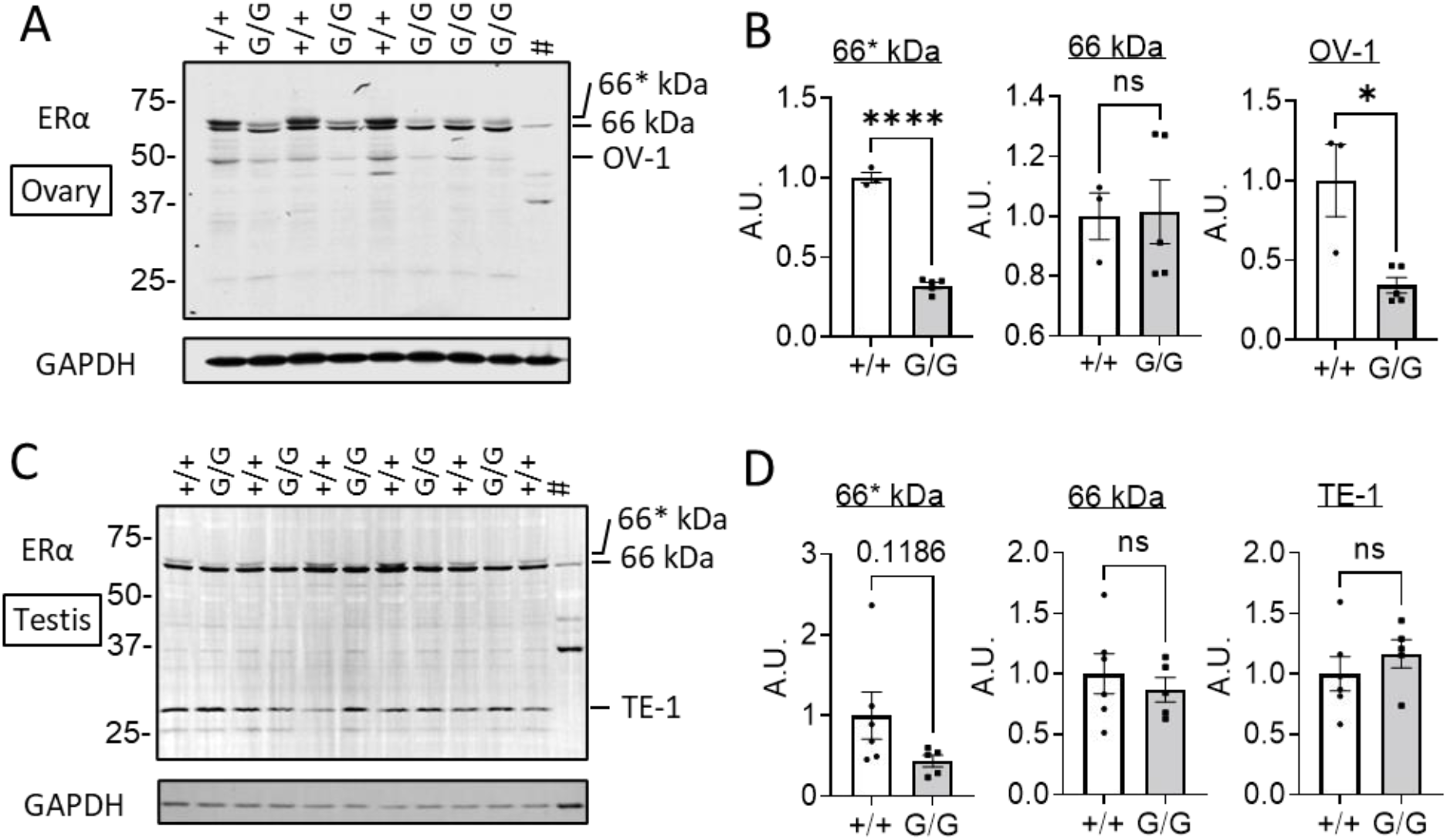
The ERα WB analysis in female ovaries and male testicles of +/+ and G/G mice. (A) ERα and GAPDH (loading control) WB images in the female ovaries (# denote a reference brain sample). (B) Quantification of 66* kDa, 66 kDa, and OV-1 bands shown in A. (C) ERα and GAPDH (loading control) WB images in the male testicles (# denote a reference brain sample). (D) Quantification of 66* kDa, 66 kDa, and TE-1 bands shown in C. The data are expressed as means ± SEM. *p < 0.05, ****p < 0.0001, by unpaired student’s t-test.

### The long ERα66* isoform is not a phosphorylated form of ERα

The mobility fluctuation around 66 kDa of ERα has been reported among the various tissues(12, 45). It has been known that the phosphorylation of ERα upshifts its mobility in SDS-PAGE(46). ERα has multiple phosphorylation sites as well as other post-translational modification sites(24). However, to our knowledge, there is no report showing that an intronic genome modification leads to a change in post-translational modifications of a targeted protein. Nonetheless, we sought to determine if ERα66* is a phosphorylated form of ERα66. Our preliminary dephosphorylation experiment with alkaline phosphatase treatment showed SDS-PAGE migration issue; the molecular weight of alkaline phosphatase (65 kDa) was too close to ERα (66 kDa) and the amount of alkaline phosphatase required for effective protein dephosphorylation disturbed ERα migration (data not shown). Therefore, we employed spontaneous dephosphorylation treatment by incubating tissue homogenates at 30°C as shown in a previous report(47). To use insulin-stimulated phosphorylation of Akt (pAKT) as a reaction control, we performed this dephosphorylation experiment in insulin-stimulated liver homogenates. First, we confirmed that 66* kDa and 66 kDa isoforms along with other 3 short molecular weight isoforms (denoted as L-1-3) are expressed in the liver of male mice (Figure 6A). The quantification revealed more than 60% reduction of 66* kDa, but not 66 kDa and L-2/3, isoform in G/G mice compared to WT littermates (Figure 6B). There was a trend toward significant reduction of L-1 band in G/G males compared to WT littermates but did not reach a statistical significance (Figure 6B). As shown in Figure 6C, incubating insulin-stimulated liver homogenates at 30°C for 3 or 6 hours was sufficient to completely eliminate pAkt, indicating the effectiveness of endogenous liver phosphatases for dephosphorylation at 30°C. Simultaneous detection of total Akt indicates that this was not due to the protein degradation, as total Akt was stable even after 6-hour incubation (Figure 6C). We then probed ERα proteins in liver homogenates incubated at 30°C for either 0 or 3 hours and found that 66* kDa band is unaffected by 3-hour incubation of liver homogenates at 30°C (Figure 6D). These results suggest that the upper 66* kDa band is not likely due to the phosphorylation of ERα66.

**Figure 6.**
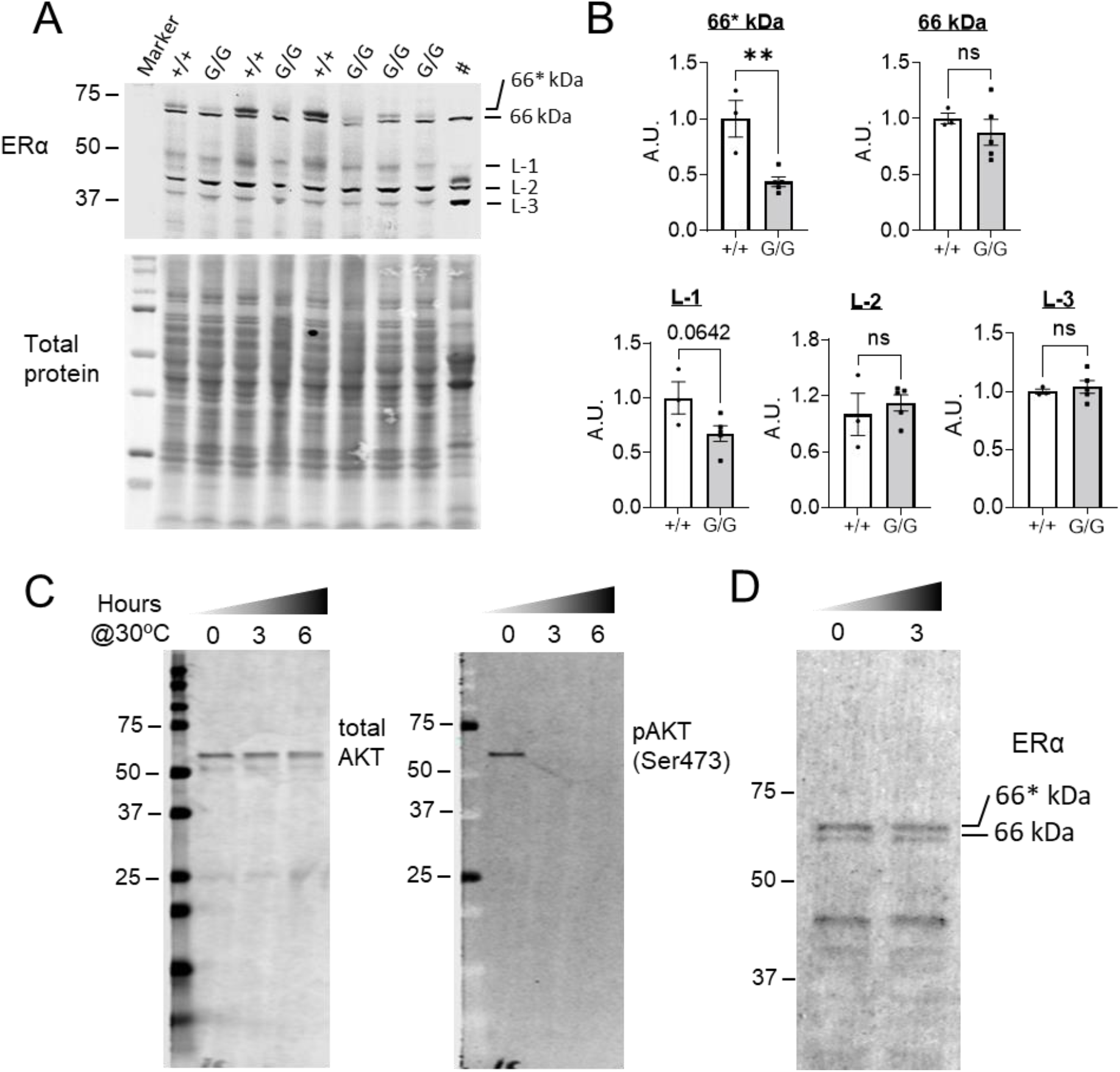
The upper 66* kDa in the liver is resistant to spontaneous dephosphorylation. (A) ERα WB image and total protein (loading control) in the liver of male mice (# denote a reference brain sample). (B) Quantification of 66* kDa, 66 kDa, and L-1-3 bands shown in A. (C) Total AKT and pAKT WB images in insulin-treated liver homogenates of WT male mice after 0, 3, and 6 hours of incubation at 30°C. (D) ERα WB image in the liver homogenates of WT male mice after 0 and 3 hours of incubation at 30°C. The data are expressed as means ± SEM. **p < 0.01 by unpaired student’s t-test.

## Discussion

Although ERα gene was cloned more than three decades ago, the list of ERα isoforms has been growing without corresponding understanding of physiological functions of each of these isoforms. In an attempt to create a Cre-restorable ERα-null mouse line, we unexpectedly proved the existence of a previously unappreciated long ERα isoform, which we termed 66* kDa isoform (ERα66*), with slightly higher molecular weight than so-called full-length 66 kDa ERα (ERα66). This long ERα66* is enriched in female reproductive organs but minimally expressed in the brain where ERα66 and other short isoforms were dominant. Strikingly, introducing a loxP-flanked TB cassette (G allele) in front of first coding exon of ERα gene resulted in selective knockdown, but not knockout, of ERα66* across the tissues and homozygous G/G mutant female mice display impaired reproductive function without significant change in body weight. These results shed light on recognizing the difference of so-called “full-length” ERα66 in the brain versus other endocrine and reproductive organs and provide new insight into the isoform-specific role of ERα in reproductive physiology and body weight homeostasis.

Varying ERα transcript variants being produced as well as the relative enrichment of those variants across the tissues perhaps contribute to various behavioral and physiological functions of ERα signaling(10-22). It has long been known that WB recognizes multiple ERα bands across the tissues; the typical ERα band appears around 66 kDa and has been considered as “full-length” ERα (ERα66) that has all reported functional domains of ERα. In addition to this full-length ERα66, two truncated isoforms with molecular weights around 36 and 46 kDa have also been well studied(48-51). ERα has been known to undergo multiple posttranslational modifications including phosphorylation, acetylation, ubiquitination, sumoylation, methylation, and palmitoylation(24), which further complicates ERα-mediated estrogen signaling. Early studies by Korach et al observed a doublet band around 66 kDa(46, 52) and the upper band was strongly induced by estradiol, preferentially extracted from the nuclear fraction, and more heavily phosphorylated(46). Joel et al reported that ERα66 was upshifted by single phosphorylation of serine 118 of ERα and downshifted by phosphatase treatment(53). Baines et al reported a doublet band around 66 kDa and they suggested the lower band was a degraded product(45). Irsik et al reported that the major band in the testis was slightly smaller than those observed in the ovaries and uterus and they suggested a possibility of a different isoform or a post-translational modification occurring in the other tissues but not in the testis(12). Additionally, Tabatadze et al reported that a doublet 66 kDa band in brain and the upper band of which was concentrated in synaptic vesicles and was prone to phosphatase treatment(54). In present study, we observed a clear doublet band around 66 kDa in all tissues examined and found that the upper ERα66* is dominant in the pituitary, ovaries, uterus, and liver, but not the brain and testis where the lower ERα66 appears to be dominant. Of note, the upper ERα66* is almost negligibly expressed in the brain of both sexes compared to other organs. Strikingly, ERα66* was significantly downregulated in G/G mice compared to control littermates in all tissues examined, while the lower ERα66 is largely unaffected. Since the genetic modification in G/G mice was made in intronic region, it is unlikely that ERα doublet formation in our samples was due to the phosphorylation of ERα66 as previously indicated (53, 54). This is further supported by our spontaneous dephosphorylation experiment in the liver homogenates where the upper ERα66* was resistant to dephosphorylation treatment. Based on these results, it is arguable that the upshift of ERα66 can occur by either an alternative splicing or different transcriptional initiation, in addition to post-translational modifications including the protein phosphorylation seen in previous studies. ERα phosphorylation was reported to be upregulated by estradiol treatment and downregulated by ovariectomy (46, 52, 53, 55), indicating that lower blood estrogens level could result in less phosphorylation of ERα. Although we did not measure blood estradiol level, G/G female mice may produce less estradiol and lead to lower ERα66* expression. However, ERα66* was dominant in the pituitary of mixed-estrus WT female mice and this was true even in male pituitary. This further indicates that the expression of ERα66* in present study is not due to ERα phosphorylation. However, we cannot fully rule out the possibility that there might be an unidentified mechanism through which a genomic *cis* element affects the process of other post-translational modifications.

In present study, we used the antibody directed against last 15 amino acid of ERα. This epitope is maintained in full-length ERα66 and ERα46 as well as some other isoforms but not in ERα36 (8). This antibody was reported to show no signal in the immunohistochemistry of ERα knockout mice brain(56). Therefore, ERα66* observed in the present study could be due to changes in the N-terminus or internal exons. Multiple non-coding exons in the 5’-UTR have been reported for ERα gene in both humans and rodents(15, 16, 22, 48). Ishii and Sakuma identified four leader exons and twelve internal exons in the 5’-UTR of mouse ERα gene (16). Although different combinations of leader exons and 5’-UTR internal exons could produce multiple splicing variants, many of them are predicted to encode identical ERα protein. However, they also suggested three combinations that could potentially produce different ERα proteins with putative 30, 34, or 35 extra amino acids on the N-terminus, which are predicted to add 3-4 kDa in molecular weight and may explain the upshift of ERα66 to ERα66*. To rigorously explore such possibility, we produced an antibody against two putative peptide antigens (GGPRAAEPSAC and CAGSQSVTRRLPLT) that encompass 24 common amino acids shared between those 3 putative extra N-terminuses. However, WB analysis using this customized antibody failed to detect a clear band around 66 kDa (data not shown). Future work should focus on determining the identify of long ERα66* isoform and how it might be different from the brain enriched ERα66 both molecularly and functionally.

Infertility or subfertility is commonly observed in various ERα mutant mouse models (1). Dupont et al reported the total absence of pregnancy in ERα KO females(5), indicating that ERα signaling is critical for normal reproductive function in female mice. Singh et al reported that conditional ERα KO female mice lacking ERα specifically in the pituitary exhibit irregular estrus cycle and subfertility(40). In the present study, we found that ERα66*, which is dominant in the pituitary, was dramatically reduced in the pituitary of G/G mice. Consistent with Singh et al, we found that G/G female mice display irregular estrus cycle (failed to enter the estrus stage) and have significantly reduced uterus weights. To our surprise, however, nearly all G/G females were capable of pregnancy in the fertility test, although they did not maintain alive newborns after the delivery. These observations imply that normal estrus cycle may not be necessary for fertilization and implantation and ERα66* in the pituitary is partially dispensable for female pregnancy. However, we cannot rule out the possibility that this was due to either residual expression of ERα66* or intact expression of ERα66 (or could be combination of both) in the pituitary of G/G mice. Nonetheless, our findings support that long ERα66* isoform is critically involved in some aspects of normal reproductive physiology in female mice.

The mechanism of subfertility (no alive pups after the delivery) observed in G/G female mice is not entirely clear. In an effort to videotape the delivery of pregnant G/G mice, we occasionally observed an alive newborn immediately after the delivery, but G/G mother seems not interested in taking care of newborn after the labor, suggesting the defect in maternal behaviors. Ogawa et al reported that conventional ERα KO female mice have impaired maternal behaviors, such as pup retrieval, and substantial number of the KO female mice also showed infanticide (57). In rodents and sheep, estrogens are known to play a key role in the onset of maternal behaviors (58) and suppression of ERα in the preoptic area (POA) abolished maternal behaviors (59). Although WB in whole brain revealed that ERα66 is dominant over ERα66* in the brain and is not affected in G/G mice, we found that this weak upper ERα66* band is also significantly decreased in G/G brains. Since we performed WB in whole brain lysate, it is possible that the upper ERα66* is specifically enriched in certain types of neurons in limited brain regions that are important for maternal behaviors, such as the POA. Another possibility of observed subfertility in G/G females is simply due to significantly reduced ERα66* across female reproductive organs and/or decreased circulating estrogen level. Since the TB cassette is flanked by loxP sites, in future studies, G/G mice can be bred to different mouse lines expressing Cre recombinase specifically in the pituitary or other female reproductive organs to fully restore hypomorphic ERα66* isoform in G/G mice to determine the sufficiency of tissue-specific role of ERα66* in female reproductive functions.

In addition to reproductive function, the role of ERα in metabolic homeostasis has been well documented. It has been shown well established that both male and female ERα KO mice gain more fat and body weight (3, 60, 61). However, in the present study, we could not detect any significant increase in body weight in young (∼10 weeks) or mature adult (∼20 weeks) G/G mice. It has been shown that metabolic effects of ERα is mainly mediated by its expression in the brain, especially in different hypothalamic nuclei(6, 7). Somewhat consistent with unaltered body weight in G/G mice, we found that, ERα66 is dominant in the brain compared to ERα66* and is largely unaffected in G/G mice. In fact, ERα66* was minimally expressed in the brain among all tissues examined in the present study. Our observations raise an interesting possibility that ERα66 and ERα66* may play dissociable roles in body weight homeostasis and reproductive function, respectively. Determination of molecular identity of ERα66* and isoform- and/or tissue-specific knockdown or knockout approaches are needed to confirm this possibility in future studies.

In conclusion, our study suggests the existence of physiologically vital ERα66* isoform that was previously mistreated as “full-length” ERα66. This long ERα66* isoform is the dominant form of ERα in HPG axis organs, but not the brain. Consistent with this, homologous G/G mutant female mice with significant knockdown of ERα66* across the tissues showed subfertility phenotype without significant increase in body weight. Although the identity and the mechanism of formation of this long ERα66* isoform remain to be determined, our study, for the first time, provides a novel insight into the isoform-specific role of ERα in female reproductive functions.

## Methods

### Targeting of the Esr1 gene using CRISPR

We used the Clustered Regularly Interspaced Short Palindromic Repeats associated protein Cas9 (CRISPR/Cas9) technology to generate a transgenic mouse line that expresses mitochondria-targeting GFP (mitoGFP) in ERα-expressing cells. Chemically modified CRISPR-Cas9 crRNA and CRISPR-Cas9 tracrRNA were purchased from IDT (Alt-R® CRISPR-Cas9 crRNA; Alt-R® CRISPR-Cas9 tracrRNA (Cat# 1072532)). The crRNA and tracrRNA were suspended in T10E0.1 and combined to 1 ug/ul (∼29.5 μM) final concentration in a 1:2 (μg: μg) ratio. The RNAs were heated at 98°C for 2 min and allowed to cool slowly to 20°C in a thermal cycler. The annealed cr:tracrRNAs were aliquoted to single-use tubes and stored at -80C. Cas9 nuclease was also purchased from IDT (Alt-R® S.p. HiFi Cas9 Nuclease). Cr:tracr:Cas9 ribonucleoprotein complexes were made by combining Cas9 protein and cr:tracrRNA in T_10_E_0.1_ (final concentrations: 500 ng/ul (∼3.1 μM) Cas9 protein and 500 ng/ul (∼14.7 μM) cr:tracrRNA). The Cas9 protein and annealed RNAs were incubated at 37°C for 10 minutes. The RNP complexes were combined with long single stranded repair template. The final concentrations of 20 ng/μl (∼0.12 μM) Cas9 protein and 20 ng/μl (∼0.6 μM) cr:tracrRNA and 10 ng/μl single stranded repair template. Pronuclear-stage embryos were collected using methods as described previously (62). Embryos were collected in KSOM media (Millipore; MR101D) and washed 3 times to remove cumulous cells. Cas9 RNPs and ds plasmid repair template were injected into the pronuclei of the collected zygotes and incubated in KSOM with amino acids at 37°C under 5% CO2 until all zygotes were injected. Fifteen to 25 embryos were immediately implanted into the oviducts of pseudo-pregnant ICR females.

Genomic DNA was prepared from the tail snip of the homozygous mice from F1 intercrosses by phenol-chloroform extraction/alcohol precipitation and dissolved in TE buffer. The targeted genomic region was amplified by PCR and the purified PCR product was processed into Sanger sequencing, which confirmed the expected genome modification.

All mice were housed in a 12-h light, 12-h dark cycle. Care of all animals and procedures were performed in accordance with protocols approved by the Institutional Animal Care and Use Committee of the University of Iowa.

### Genotyping of mice

Genomic DNA from tail or distal portion of digit were used as templates in genotyping PCR using primers P1 (5’-GTGCACGCACAGTGTGATGTTCC-3’), P2 (5’-CGCTCCCGGGTTCTCCAACTTTA-3’), and P3 (5’-TTCTACCCCAGACCTTGGGACCAC-3’), which produce 122-bp band as WT allele, 307-bp band as mutant allele, and 197-bp band in case that Cre-mediated recombination occurs.

### Body weight measurement

After weaning till 10 weeks of age, both male and female wild type and mutant mice were weighed every four days right before dark cycle onset to examine the effect of ERα knockdown on body weight.

### Estrus cycles

Starting from 6-7 weeks of age, estrus cycle of WT and G/G mutant female mice were checked daily for 2-3 weeks. Vaginal smears were collected every early afternoon. Estrus cycle stages were determined based on the criteria in Caligioni (63).

### Fertility test

For evaluation of female fertility, fertility-proven WT males were placed with 4- to 6-week-old wild-type, +/G, and G/G females as in Krege et al (64). Once pregnancy was confirmed by belly expansion, females were singly housed and cages were monitored daily for the delivery.

### Western blot

Nine- to 18-week-old wild-type and G/G mice were sacrificed by cervical dislocation and the tissues were comprehensively collected, immediately snap-frozen in liquid nitrogen, and stored at -80°C until use. Frozen tissues were homogenized with a sonicator in ice-cold radioimmunoprecipitation assay lysis buffer (Thermo Fisher Scientific, Waltham, MA) containing Complete protease inhibitor cocktail (Millipore Sigma, Burlington, MA). Protein concentration was determined using Pierce BCA Protein Assay Kit (Thermo Fisher Scientific). Equal amounts of protein (13.5 to 30 µg) per tissue were separated by 4% to 20% sodium dodecyl sulfate– polyacrylamide gel electrophoresis gradient gels electrophoresis (SDS-PAGE) and transferred to a polyvinylidene fluoride (PVDF) membrane by semi-dry electrotranfer method (Trans-Blot Turbo Blotting System, Bio-Rad, Hercules, CA). After the transfer, the entire blots were stained with Revert Total Protein Stain Kit (Li-Cor, Lincoln, NE) for the quantification of total protein on the blots. After rinses in Tris Buffered Saline containing 0.1% Tween-20 (TBSTw), the blots were blocked in blocking buffer (TBSTw containing 5% skim milk) for one hour at room temperature while gentle agitation followed by the incubation in blocking buffer containing ERα primary antibody (Millipore Sigma, Burlington, MA, cat#06-935, 1:1000) overnight at 4°C while gentle agitation. The signals were detected with Li-Cor IRDye secondary antibody (Li-Cor, 1:15000) and visualized using Li-Cor Odyssey infrared scanner (Li-Cor, Lincoln, NE) or Azure Biomolecular Imager (Azure Biosystems, Dublin, CA). Signal intensity was measured and analyzed by Bio-Rad Image Lab. For each experiment, signal intensity of targeted proteins was normalized to GAPDH expression levels or total protein signal intensity and then compared between the groups.

### Dephosphorylation treatment

The spontaneous dephosphorylation of the protein homogenates was induced as described in Bedet et al. with slight modification(47). Organs were dissected and sonicated in ice-cold lysis buffer (10 mM Tris-HCl, 10 mM NaCl, 1% Nonidet P-40, pH8.0) supplemented with 1X Protease Inhibitor Cocktail III (Research Products International, Mt. Prospect, IL) until the homogenate was easily pipetted without clogging. After clearing by centrifugation, the protein concentration was determined using Pierce BCA Protein Assay Kit (Thermo Fisher Scientific). Tissue homogenates were diluted in the above lysis buffer to adjust the protein concentration (2-4 mg/ml). Then the homogenates were diluted 10 times with the dephosphorylation buffer (50 mM Tris-HCl, pH7.4, 5 mM DTT, 0.05% SDS, protease inhibitor cocktail) and incubated at 30°C for 0, 3, or 6 hours. Samples were then mixed with Laemmli sample buffer as having same protein concentration determined earlier, boiled for 5 min at 95°C, and stored at -80°C. The mobility of ERα protein was then evaluated in Western blot analysis as described above.

### Histology

Both male and female mutant mice (8–10 weeks of age) were anesthetized with isoflurane inhalation and were perfused intracardially with cold saline followed by 10% formalin. The brains and pituitaries were removed, post-fixed overnight in the same fixative diluted half with 20% sucrose, and then immersed in 20% sucrose in PBS at 4 °C until sinking in the solution. 30-μm-thick coronal sections throughout whole brains were prepared by freezing microtome, collected in cryoprotective solution (20% glycerol, 30% ethylene glycol in PBS) and stored at −20 °C until use. Brain sections were thoroughly rinsed in PBS, incubated in blocking buffer (PBS containing 5% normal donkey serum (Jackson ImmunoResearch Laboratories, West Grove, PA, cat#017-000-121) and 0.05% Triton X-100) for 1 hour at room temperature, and then incubated in blocking buffer with ERα antibody (Millipore Sigma, Burlington, MA, cat#06-935, 1:1000) overnight at 4°C. On the following day sections were rinsed three exchanges of PBS for 10 min each and incubated with secondary antibody in blocking buffer for two hours in the dark at room temperature. The stained sections were rinsed in three exchanges of PBS for 10 min each, mounted on SuperFrost Plus glass slides (Fisher Scientific, Waltham, MA, cat#12-550-15), and cover-slipped with Vectashield hardset antifade mounting medium with DAPI (Vector Laboratories, Burlington, CA, cat#H-1500). Fluorescent images were captured with Olympus VS120 slide scanner microscope or Olympus FV3000 confocal microscope.

For the histological examination of female organs, mice were transcardially perfused with 10% formalin and female organs including ovaries, uterus, and vagina, were dissected together and sent to the UI Comparative Pathology Laboratory for paraffin embedding, sectioning, and hematoxylin and eosin staining.

### Statistics

All values were expressed as mean ± SEM. Statistical analyses were performed using GraphPad Prism. ERα expression in Western blot analysis were compared with Student’s t-test. The body weight changes were compared with one-way analysis of variance (ANOVA). P <0.05 was considered to be statistically significant.

## Acknowledgement

Transgenic mice were generated at the University of Iowa Genome Editing Core Facility directed by William J. Paradee, PhD, and supported in part by grants from the National Institutes of Health and from the Roy J. and Lucille A. Carver College of Medicine. This work was supported by grants from the National Institutes of Health (HL127673 and HL084207 to HC). We wish to thank Norma Sinclair, Patricia Yarolem, Joanne Schwarting and Rongbin Guan for their technical expertise in generating transgenic mice. We also acknowledge the expert assistance of Katherine N. Gibson-Corley in the UI Comparative Pathology Laboratory.

## Author Contributions

KS and HC designed the study. KS, JED, SRR, and MJK performed data acquisition, and BAT, US, GD, DY, and JJ assisted in production of study mice. KS and HC wrote the manuscript, and US, GD, DY, and JJ revised the manuscript. HC is the guarantor of this work and, as such, had full access to all the data in this study.

## Figures and legends

**Supplemental table S1.** Frequency of sex and genotypes of offspring from ERα^+/G^ intercrosses

|  | +/+ | +/G | G/G |
| --- | --- | --- | --- |
| Male | 21 | 29 | 16 |
| female | 28 | 37 | 20 |
| Total | 49 (32.5%) | 66 (43.7%) | 36 (23.8%) |

Numbers of offspring upon weaning at postnatal day 18-24.

**Supplemental figure S1.**
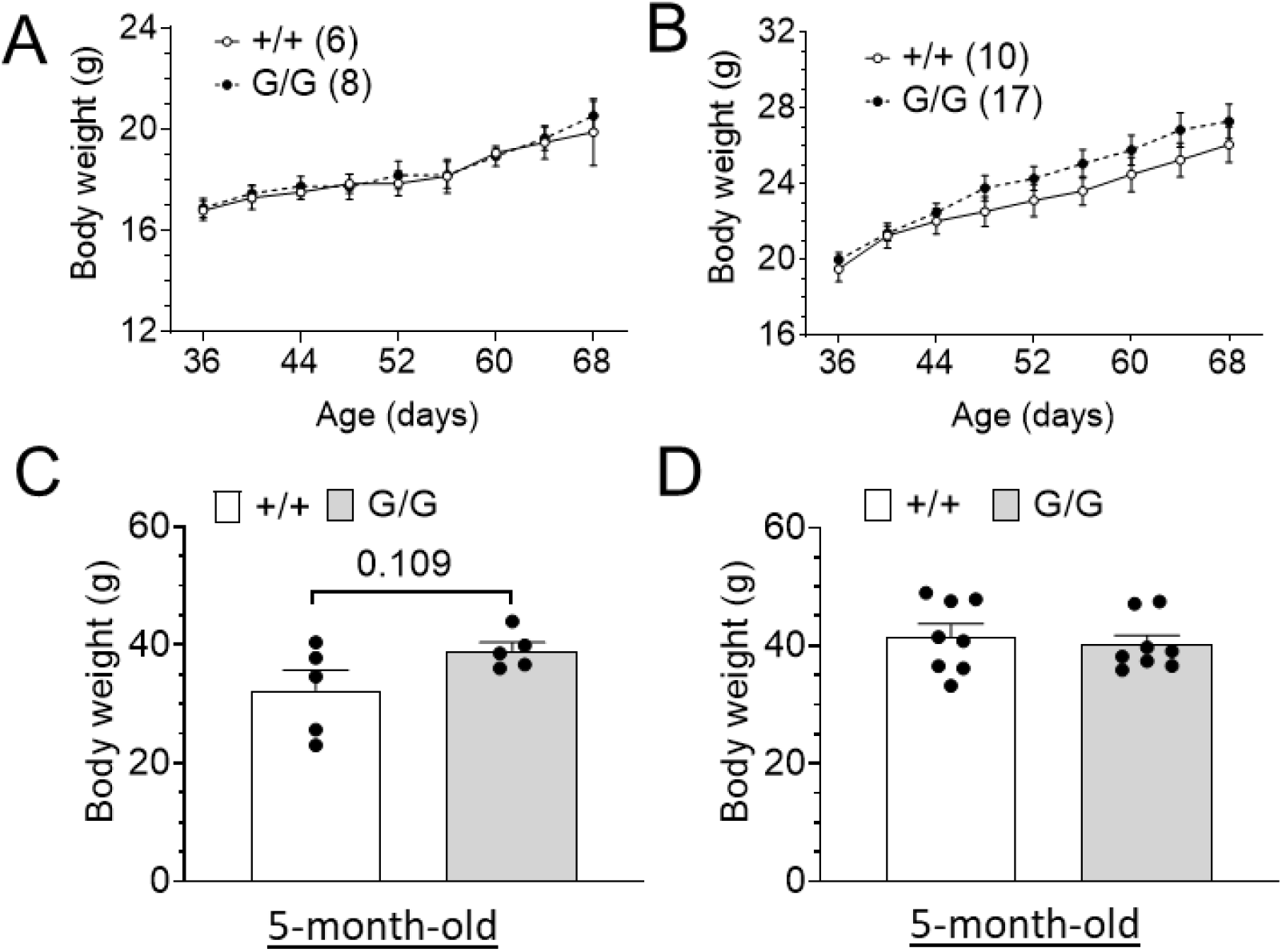
Body weight gain in +/+ and G/G mice. (A, B) Body weight growth curve in young female (A) and male (B) mice. (C, D) Body weight in 5-month-old male (C) and female (D) mice. The data are expressed as means ± SEM.

**Supplemental figure S2.**
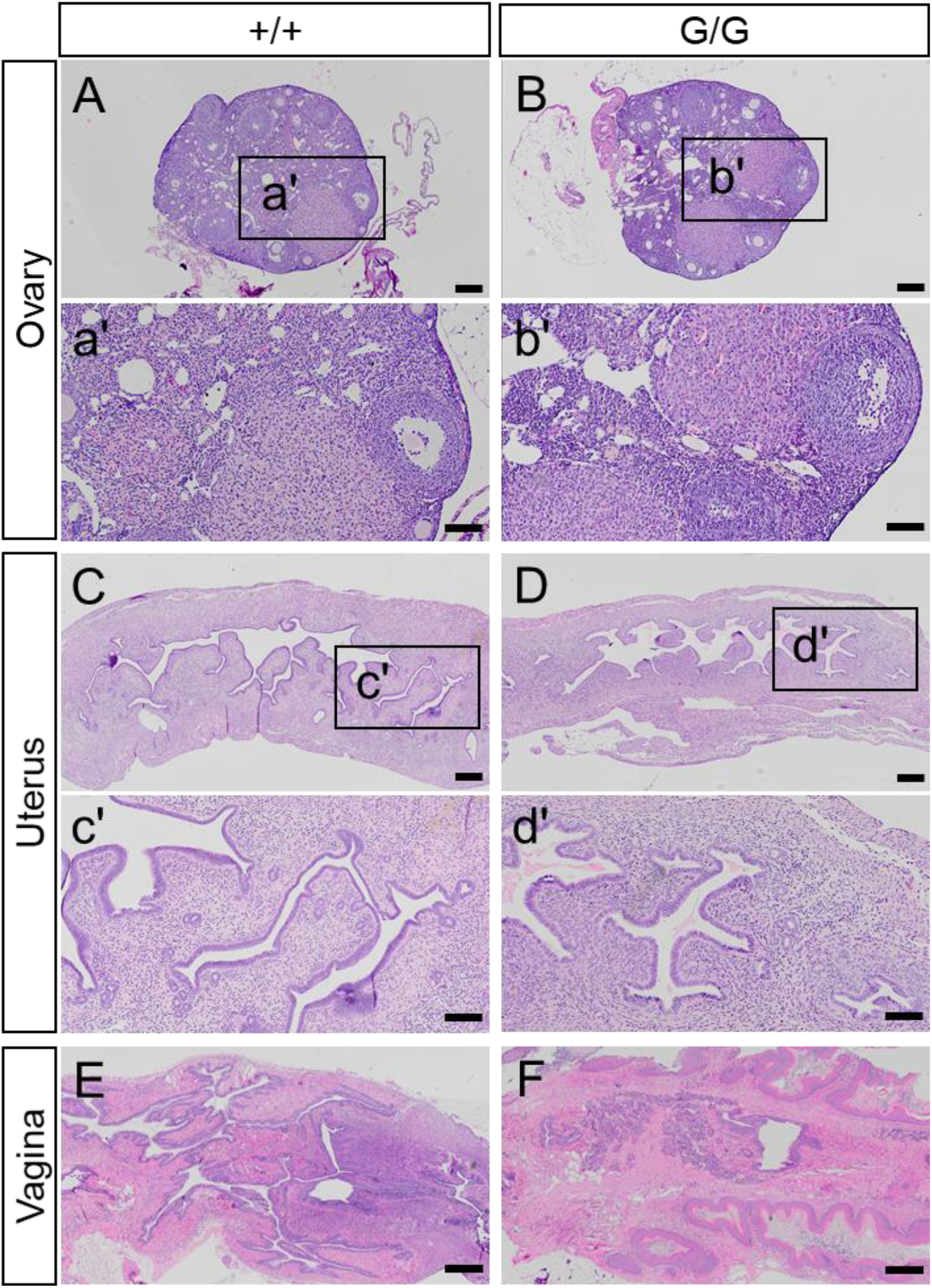
Representative HE staining images showing intact gross anatomy in the ovary (A, B), the uterus (C, D), and the vagina (E, F) of G/G female mice. Scale bar: 200 µm for A-D; 500 µm for E and F; 100 µm for a’-d’.

## References

1. Hewitt SC, and Korach KS. Estrogen receptors: New directions in the new millennium. Endocrine Reviews. 2018;39(5):664–75.

2. Morselli E, Santos RS, Criollo A, Nelson MD, Palmer BF, and Clegg DJ. The effects of oestrogens and their receptors on cardiometabolic health. Nat Rev Endocrinol. 2017;13(6):352–64.

3. Heine PA, Taylor JA, Iwamoto GA, Lubahn DB, and Cooke PS. Increased adipose tissue in male and female estrogen receptor-alpha knockout mice. Proc Natl Acad Sci U S A. 2000;97(23):12729–34.

4. Lubahn DB, Moyer JS, Golding TS, Couse JF, Korach KS, and Smithies O. Alteration of reproductive function but not prenatal sexual development after insertional disruption of the mouse estrogen receptor gene. Proc Natl Acad Sci U S A. 1993;90(23):11162–6.

5. Dupont S, Krust A, Gansmuller A, Dierich A, Chambon P, and Mark M. Effect of single and compound knockouts of estrogen receptors alpha (ERalpha) and beta (ERbeta) on mouse reproductive phenotypes. Development. 2000;127(19):4277–91.

6. Musatov S, Chen W, Pfaff DW, Mobbs CV, Yang X-J, Clegg DJ, et al. Silencing of estrogen receptor alpha in the ventromedial nucleus of hypothalamus leads to metabolic syndrome. Proceedings of the National Academy of Sciences of the United States of America. 2007;104(7):2501–6.

7. Xu Y, Nedungadi TP, Zhu L, Sobhani N, Irani BG, Davis KE, et al. Distinct hypothalamic neurons mediate estrogenic effects on energy homeostasis and reproduction. Cell Metabolism. 2011;14(4):453–65.

8. Taylor SE, Martin-Hirsch PL, and Martin FL. Oestrogen receptor splice variants in the pathogenesis of disease. Cancer Lett. 2010;288(2):133–48.

9. Kundu P, Li M, Lu R, Stefani E, and Toro L. Regulation of transcriptional activation function of rat estrogen receptor alpha (ERalpha) by novel C-terminal splice inserts. Mol Cell Endocrinol. 2015;401:202–12.

10. Flouriot G, Griffin C, Kenealy M, Sonntag-Buck V, and Gannon F. Differentially expressed messenger RNA isoforms of the human estrogen receptor-alpha gene are generated by alternative splicing and promoter usage. Molecular endocrinology (Baltimore, Md). 1998;12(12):1939–54.

11. Hattori Y, Ishii H, Morita A, Sakuma Y, and Ozawa H. Characterization of the fundamental properties of the N-terminal truncation (δ exon 1) variant of estrogen receptor α in the rat. Gene. 2015;571(1):117–25.

12. Irsik DL, Carmines PK, and Lane PH. Classical estrogen receptors and ERα splice variants in the mouse. PloS one. 2013;8(8):e70926–e.

13. Ishii H, Hattori Y, Munetomo A, Watanabe H, Sakuma Y, and Ozawa H. Characterization of rodent constitutively active estrogen receptor α variants and their constitutive transactivation mechanisms. General and comparative endocrinology. 2017;248:16–26.

14. Ishii H, Kobayashi M, Munetomo A, Miyamoto T, and Sakuma Y. Novel splicing events and post-transcriptional regulation of human estrogen receptor α e isoforms. Journal of Steroid Biochemistry and Molecular Biology. 2013;133(1):120–8.

15. Ishii H, Kobayashi M, and Sakuma Y. Alternative promoter usage and alternative splicing of the rat estrogen receptor α gene generate numerous mRNA variants with distinct 5′-ends. Journal of Steroid Biochemistry and Molecular Biology. 2010;118(1-2):59–69.

16. Ishii H, and Sakuma Y. Complex organization of the 5 ′ -untranslated region of the mouse estrogen receptor α gene: Identification of numerous mRNA transcripts with distinct 5′-ends. Journal of Steroid Biochemistry and Molecular Biology. 2011;125(3-5):211–8.

17. Ishunina TA, Sluiter AA, Swaab DF, and Verwer RWH. Transcriptional activity of human brain estrogen receptor-α splice variants: evidence for cell type-specific regulation. Brain research. 2013;1500:1–9.

18. Koš M, Denger S, Reid G, and Gannon F. Upstream open reading frames regulate the translation of the multiple mRNA variants of the estrogen receptor α. Journal of Biological Chemistry. 2002;277(40):37131–8.

19. Kos M, O’Brien S, Flouriot G, and Gannon F. Tissue-specific expression of multiple mRNA variants of the mouse estrogen receptor alpha gene. FEBS letters. 2000;477(1-2):15–20.

20. Kos M, Reid G, Denger S, and Gannon F. Minireview: genomic organization of the human ERalpha gene promoter region. Molecular endocrinology (Baltimore, Md). 2001;15(12):2057–63.

21. Swope DL, Harrell JC, Mahato D, and Korach KS. Genomic structure and identification of a truncated variant message of the mouse estrogen receptor alpha gene. Gene. 2002;294(1-2):239–47.

22. Kobayashi M, Ishii H, and Sakuma Y. Identification of novel splicing events and post-transcriptional regulation of human estrogen receptor alpha F isoforms. Mol Cell Endocrinol. 2011;333(1):55–61.

23. Sun JW, Collins JM, Ling D, and Wang D. Highly Variable Expression of ESR1 Splice Variants in Human Liver: Implication in the Liver Gene Expression Regulation and Inter-Person Variability in Drug Metabolism and Liver Related Diseases. J Mol Genet Med. 2019;13(3).

24. Le Romancer M, Poulard C, Cohen P, Sentis SP, Renoir JM, and Corbo L. Cracking the estrogen receptor’s posttranslational code in breast tumors. Endocrine Reviews. 2011;32(5):597–622.

25. Anbalagan M, and Rowan BG. Estrogen receptor alpha phosphorylation and its functional impact in human breast cancer. Mol Cell Endocrinol. 2015;418 Pt 3p264-72.

26. Ohlsson C, Gustafsson KL, Farman HH, Henning P, Lionikaite V, Moverare-Skrtic S, et al. Phosphorylation site S122 in estrogen receptor alpha has a tissue-dependent role in female mice. FASEB J. 2020;34(12):15991–6002.

27. Adlanmerini M, Solinhac R, Abot A, Fabre A, Raymond-Letron I, Guihot AL, et al. Mutation of the palmitoylation site of estrogen receptor alpha in vivo reveals tissue-specific roles for membrane versus nuclear actions. Proc Natl Acad Sci U S A. 2014;111(2):E283–90.

28. Fugger HN, Foster TC, Gustafsson J, and Rissman EF. Novel effects of estradiol and estrogen receptor alpha and beta on cognitive function. Brain Res. 2000;883(2):258–64.

29. Li L, Fan X, Warner M, Xu XJ, Gustafsson JA, and Wiesenfeld-Hallin Z. Ablation of estrogen receptor alpha or beta eliminates sex differences in mechanical pain threshold in normal and inflamed mice. Pain. 2009;143(1-2):37–40.

30. Berglund ED, Vianna CR, Donato J, Kim MH, Chuang JC, Lee CE, et al. Direct leptin action on POMC neurons regulates glucose homeostasis and hepatic insulin sensitivity in mice. Journal of Clinical Investigation. 2012;122(3):1000–9.

31. Balthasar N, Dalgaard LT, Lee CE, Yu J, Funahashi H, Williams T, et al. Divergence of melanocortin pathways in the control of food intake and energy expenditure. Cell. 2005;123(3):493–505.

32. Zigman JM, Nakano Y, Coppari R, Balthasar N, Marcus JN, Lee CE, et al. Mice lacking ghrelin receptors resist the development of diet-induced obesity. Journal of Clinical Investigation. 2005;115(12):3564–72.

33. Xu Y, Jones JE, Kohno D, Williams KW, Lee CE, Choi MJ, et al. 5-HT2CRs Expressed by Pro-Opiomelanocortin Neurons Regulate Energy Homeostasis. Neuron. 2008;60(4):582–9.

34. Klinge CM. Estrogenic control of mitochondrial function and biogenesis. J Cell Biochem. 2008;105(6):1342–51.

35. Ventura-Clapier R, Piquereau J, Veksler V, and Garnier A. Estrogens, Estrogen Receptors Effects on Cardiac and Skeletal Muscle Mitochondria. Front Endocrinol (Lausanne). 2019;10:557.

36. Saito K, He Y, Yan X, Yang Y, Wang C, Xu P, et al. Visualizing estrogen receptor-α-expressing neurons using a new ERα-ZsGreen reporter mouse line. Metabolism: Clinical and Experimental. 2016;65(4):522–32.

37. Merchenthaler I, Lane MV, Numan S, and Dellovade TL. Distribution of estrogen receptor alpha and beta in the mouse central nervous system: in vivo autoradiographic and immunocytochemical analyses. The Journal of comparative neurology. 2004;473(2):270–91.

38. Arao Y, Hamilton KJ, Wu SP, Tsai MJ, DeMayo FJ, and Korach KS. Dysregulation of hypothalamic-pituitary estrogen receptor alpha-mediated signaling causes episodic LH secretion and cystic ovary. FASEB J. 2019;33(6):7375–86.

39. Gieske MC, Kim HJ, Legan SJ, Koo Y, Krust A, Chambon P, et al. Pituitary gonadotroph estrogen receptor-alpha is necessary for fertility in females. Endocrinology. 2008;149(1):20–7.

40. Singh SP, Wolfe A, Ng Y, DiVall SA, Buggs C, Levine JE, et al. Impaired estrogen feedback and infertility in female mice with pituitary-specific deletion of estrogen receptor alpha (ESR1). Biology of reproduction. 2009;81(3):488–96.

41. Antonson P, Omoto Y, Humire P, and Gustafsson JÅ. Generation of ERα-floxed and knockout mice using the Cre/LoxP system. Biochemical and biophysical research communications. 2012;424(4):710–6.

42. Couse JF, Hewitt SC, Bunch DO, Sar M, Walker VR, Davis BJ, et al. Postnatal sex reversal of the ovaries in mice lacking estrogen receptors alpha and beta. Science. 1999;286(5448):2328–31.

43. Mahboobifard F, Bidari-Zerehpoosh F, Davoudi Z, Panahi M, Dargahi L, Pourgholami MH, et al. Expression patterns of ERalpha66 and its novel variant isoform ERalpha36 in lactotroph pituitary adenomas and associations with clinicopathological characteristics. Pituitary. 2020;23(3):232–45.

44. Yan Y, Yu L, Castro L, and Dixon D. ERalpha36, a variant of estrogen receptor alpha, is predominantly localized in mitochondria of human uterine smooth muscle and leiomyoma cells. PLoS One. 2017;12(10):e0186078.

45. Baines H, Nwagwu MO, Hastie GR, Wiles RA, Mayhew TM, and Ebling FJP. Effects of estradiol and FSH on maturation of the testis in the hypogonadal (hpg) mouse. Reproductive Biology and Endocrinology. 2008;6:1–10.

46. Washburn T, Hocutt A, Brautigan DL, and Korach KS. Uterine estrogen receptor in vivo: phosphorylation of nuclear specific forms on serine residues. Molecular endocrinology (Baltimore, Md). 1991;5(2):235–42.

47. Bedet C, Isambert MF, Henry JP, and Gasnier B. Constitutive phosphorylation of the vesicular inhibitory amino acid transporter in rat central nervous system. J Neurochem. 2000;75(4):1654–63.

48. Flouriot G, Brand H, Denger S, Metivier R, Kos M, Reid G, et al. Identification of a new isoform of the human estrogen receptor-alpha (hER-alpha) that is encoded by distinct transcripts and that is able to repress hER-alpha activation function 1. The EMBO journal. 2000;19(17):4688–700.

49. Zhao YW, Zhang X, Shen P, Loggie BW, Chang Y, and Deuel TF. Identification, cloning, and expression of human estrogen receptor-α36, a novel variant of human estrogen receptor-α66. Biochemical and Biophysical Research Communications. 2005;336(4):1023–7.

50. Wang Z, Zhang X, Shen P, Loggie BW, Chang Y, and Deuel TF. A variant of estrogen receptor-{alpha}, hER-{alpha}36: transduction of estrogen- and antiestrogen-dependent membrane-initiated mitogenic signaling. Proceedings of the National Academy of Sciences of the United States of America. 2006;103(24):9063–8.

51. Zhang XT, Kang LG, Ding L, Vranic S, Gatalica Z, and Wang ZY. A positive feedback loop of ER-α36/EGFR promotes malignant growth of ER-negative breast cancer cells. Oncogene. 2011;30(7):770–80.

52. Golding TS, and Korach KS. Nuclear estrogen receptor molecular heterogeneity in the mouse uterus. Proceedings of the National Academy of Sciences of the United States of America. 1988;85(1):69–73.

53. Joel PB, Traish AM, and Lannigan DA. Estradiol-induced phosphorylation of serine 118 in the estrogen receptor is independent of p42/p44 mitogen-activated protein kinase. The Journal of biological chemistry. 1998;273(21):13317–23.

54. Tabatadze N, Smejkalova T, and Woolley CS. Distribution and posttranslational modification of synaptic ERalpha in the adult female rat hippocampus. Endocrinology. 2013;154(2):819–30.

55. Stirone C, Duckles SP, and Krause DN. Multiple forms of estrogen receptor-alpha in cerebral blood vessels: regulation by estrogen. American journal of physiology Endocrinology and metabolism. 2003;284(1):E184–92.

56. Vanderhorst VG, Gustafsson JA, and Ulfhake B. Estrogen receptor-alpha and -beta immunoreactive neurons in the brainstem and spinal cord of male and female mice: relationships to monoaminergic, cholinergic, and spinal projection systems. J Comp Neurol. 2005;488(2):152–79.

57. Ogawa S, Eng V, Taylor J, Lubahn DB, Korach KS, and Pfaff DW. Roles of Estrogen Receptor-α Gene Expression in Reproduction-Related Behaviors in Female Mice. Endocrinology. 1998;139(12):5070–81.

58. Bridges RS. Neuroendocrine regulation of maternal behavior. Frontiers in neuroendocrinology. 2015;36:178–96.

59. Ribeiro AC, Musatov S, Shteyler A, Simanduyev S, Arrieta-Cruz I, Ogawa S, et al. siRNA silencing of estrogen receptor-α expression specifically in medial preoptic area neurons abolishes maternal care in female mice. Proceedings of the National Academy of Sciences of the United States of America. 2012;109(40):16324–9.

60. Ohlsson C, Hellberg N, Parini P, Vidal O, Bohlooly YM, Rudling M, et al. Obesity and disturbed lipoprotein profile in estrogen receptor-alpha-deficient male mice. Biochem Biophys Res Commun. 2000;278(3):640–5.

61. Vidal O, Lindberg M, Savendahl L, Lubahn DB, Ritzen EM, Gustafsson JA, et al. Disproportional body growth in female estrogen receptor-alpha-inactivated mice. Biochem Biophys Res Commun. 1999;265(2):569–71.

62. Pinkert CA. Transgenic animal technology : a laboratory handbook. Academic Press; 2002.

63. Caligioni CS. 2009:Appendix 4I-Appendix I.

64. Krege JH, Hodgin JB, Couse JF, Enmark E, Warner M, Mahler JF, et al. Generation and reproductive phenotypes of mice lacking estrogen receptor beta. Proc Natl Acad Sci U S A. 1998;95(26):15677–82.

